# Attenuation of obesity and related metabolic disorders by the individual or combination treatment with IL-2/anti-IL-2 complex and hyperbaric oxygen

**DOI:** 10.1101/351841

**Authors:** Eun-Jeong Choi, Hyung-Ran Kim, Kie Jeong-Hae, Byung-In Moon, Ju-Young Seoh

## Abstract

Obesity is the disease accumulating excessive fat in the body. The prevalence of obesity and related metabolic disorders is increasing every year worldwide. Immunologically, obesity is a chronic low-grade inflammatory state with the increase of M1 macrophages and decrease of regulatory T cells (Tregs). IL-2/anti-IL-2 complex (IL-2C) and hyperbaric oxygen (HBO) are known to expand Tregs *in vivo* and suppress inflammation. Therefore, in this study, IL-2C and HBO were investigated for the preventive effect of obesity and related metabolic disorders. Male C57BL/6 mice were fed with a high-fat diet (HFD) for 16 weeks, and counterparts were fed with a low-fat diet (LFD). At the end of the experiment, the body weight gain and impaired glucose metabolism, elevated levels of insulin and total cholesterol induced by HFD were improved by the individual or combination treatment with Il-2C and HBO. Histological examination of the epididymal white adipose tissue showed adipocyte hypertrophy and many crown-like structures in the HFD control groups. In addition, the liver showed the progression of non-alcoholic fatty liver disease (NAFLD) in the HFD control groups, but it was significantly improved by the individual or combination treatment with IL-2C and HBO.

As for the underlying mechanism, inflammation induced by obesity was decreased, and HIF-1α expression by adipocyte hypertrophy was also reduced by the individual or combination treatment with IL-2C and HBO. In addition, adipose tissue browning was activated in brown and inguinal adipose tissue, and the expression of UCP-1 involved in the thermogenesis was increased by the individual or combination treatment with IL-2C and HBO. Overall, these results suggested that IL-2C and HBO might be a new promising immunotherapy for the treatment of obesity and related metabolic disorders by regulation of inflammation and activation of adipose tissue browning.

## Introduction

Obesity comes from an imbalance between energy uptake and consumption [1]. It is the state of excessive fat accumulated in the body and causes many adverse effects on the health. WHO reported more than 1.9 billion adults were overweight, and out of these over 650 million were obese in 2016 [2]. Nowadays, this issue becomes more serious as obesity is becoming more prevalent in children and adolescents [3]. Obesity is associated with diverse metabolic disorders, including type 2 diabetes [4] and non-alcoholic fatty liver disease (NAFLD) [5], vascular diseases including atherosclerosis [6] and cardiovascular diseases [7], and several cancers [8]. Ultimately, the obesity shortens lifespan. For the treatment of obesity, the most effective and safe method is changing the dietary regime and lifestyle [9], but it is not easy. In serious cases, medical and even surgical interventions are considered [10]. However, there are many side effects and complications [11, 12], and thus, it is still necessary to develop safer alternative methods for the treatment of obesity.

Adipose tissue is important for the maintenance of life [13]. In the past, it was simply thought of as an organ storing energy as a fat to prepare for the future starvation. Nowadays, it is considered as an
endocrine organ capable to mediate biological effects on the metabolism and inflammation [14]. Adipose tissues release adipokines such as various hormones and cytokines that control energy balance by regulating appetitive signals and metabolic activity [15]. On the other hand, adipose tissue undergoes dynamic changes in response to stimuli such as the nutritional status and temperature change. These stimuli induce remodeling of the adipose tissue as well as change the size and number of adipocytes [16]. There are two types of adipose tissues, white adipose tissue (WAT) and brown adipose tissue (BAT) [17, 18]. Both the WAT and BAT are involved in energy balance, but they are distinct in the anatomical locations, morphology, functions, and regulations. Especially, uncoupling protein (UCP-1) is mainly expressed in the BAT and play an important role in the metabolic and energy balance, such as cold- or diet-induced non-shivering thermogenesis [19]. Recently, the brown-like adipocytes were discovered in the WAT and it was called as ‘beige or brite’ adipocytes [20]. It implies the increase in metabolic activity, and therefore, browning of WAT may be a new strategy of anti-obesity therapy [21]. As obesity progresses, adipose tissues exhibit various changes. Lipid metabolites in the adipose tissues are accumulated by incomplete β-oxidation and increase of esterification [22, 23]. Adipocyte hypertrophy and the decrease of blood flow induce hypoxic condition in the adipose tissues and activation of the hypoxia-inducible factor-1 alpha (HIF-1α). In addition, infiltration of inflammatory cells including adipose tissue macrophages (ATMs) and formation of crown-like structures around dead adipocytes were observed [24], implying chronic low-grade inflammation [25]. ATMs exist in the visceral adipose tissue (VAT) and other metabolic tissues (liver, skeletal muscles) and secrete pro-inflammatory cytokines such as IL-1β, IL-6 and TNF-α [26]. These circulating cytokines result in insulin resistance and play a critical role in the metabolic dysfunction. In addition, obesity induces the conversion of M2 (or alternatively activated) macrophages into M1 (or classically activated) macrophages. On the other hand, regulatory T cells (Tregs), that play a critical role in the maintenance of immune homeostasis *in vivo,* are decreased in obese adipose tissues. Thus, it could be hypothesized that it can be reverse the altered composition of inflammatory cells by increasing the Tregs as well as decreasing the M1 macrophages by the individual or combination treatment with IL-2/anti-IL-2 complex (IL-2C) and HBO. IL-2 is an important cytokine for the survival and function of Tregs and *in vivo* injection of IL-2 can induce expansion of Tregs [27]. Meanwhile, the half-life of IL-2 is very short and is rapidly removed from the circulation via renal clearance [28]. Therefore, the IL-2 coupled with an anti-IL-2 monoclonal antibody (JES6-1A12) is a very effective method for the selective expansion of CD4^+^Foxp3^+^ Tregs. It might be related with masking of motifs binding with IL-2Rβ on cytotoxic T cells and NK cells as well as prolongation of half-life. IL-2C has been shown to suppress several autoimmune or inflammatory diseases through expanding CD4+Foxp3+ Tregs [29–31]. Hyperbaric oxygen (HBO) is another way to expand CD4^+^ Foxp3^+^ Tregs [32, 33]. HBO is breathing of 100% oxygen with increased atmospheric pressure (2 - 3 ATA). During the treatment, the arterial oxygen tension reached to almost 2,000 mmHg, and in the tissue about 200 to 400 mmHg [34]. It was initially used for the treatment of arterial gas embolism [35] and decompression sickness. Nowadays, HBO is applied to the diverse diseases, and it has improved diseases and symptoms. Breathing greater than 1 ATA of oxygen can increase of reactive oxygen species (ROS) level in the tissue [36]. Slight transient elevation of ROS increases the number and function of Tregs and suppresses inflammation. These effects have been reported that HBO attenuated autoimmune diseases such as atopic dermatitis [37] and psoriasis [38] by increasing the number and function of Tregs *in vivo.* In addition, it has been also reported that HBO treatment attenuated obesity in an animal model and improved altered glucose metabolism in obese and type 2 diabetic people [39–41].

Thus, both the IL-2C and HBO may expand Tregs and suppress chronic low-grade inflammatory status in obesity, and synergy can be anticipated. Therefore, in the present study, the individual or combination treatment of IL-2C and HBO were investigated for the preventive effects on obesity and related metabolic disorders induced by a high-fat diet.

## Materials and methods

### Mice

This study was approved by the Institutional Animal Care and Use Committee of Ewha Womans University Graduate School of Medicine (IACUC approval number: 14-0263). Mice (C57BL/6, male, 6 weeks old) were purchased from Central Lab Animal Inc. (Seoul, Korea). Mice were housed in a specific pathogen-free facility at Ewha Womans University. For animal care, a room was controlled with a 12 hours light/dark cycle, 50% humidity and *ad libitum* access to food and water treated with γ-irradiation.

### Animal experiment

Seven-week-old C57BL/6J male mice were used in the experiment. Mice were fed with a high-fat diet (HFD, 60 kcal% fat diets, 5.24 kcal/g, D12492, Research Diets, New Brunswick, NJ, USA) or a low-fat diet (LFD, 10 kcal% fat diets, 3.85 kcal/g, D12450B, Research Diets) for 16 weeks. The specific aim of this experiment is to investigate if IL-2C and/or hyperbaric oxygen (HBO), that are known to expand Tregs *in vivo*, attenuate HFD-induced obesity and related metabolic disorders.

Accordingly, the mice in the LFD and HFD groups were further divided into 5 sub-groups treated with IL-2C, HBO or both, and the control groups treated with none or PBS (vehicle for IL-2C). Each sub-group was composed of 8 - 16 mice. Body weights and food intakes were measured every week during the experiment. The IL-2C mixture was prepared by mixing mouse recombinant (r) IL-2 (eBioscience, San Diego, CA, USA) and anti-mouse IL-2 IgG (JES6-1A12, eBioscience) at a ratio of 1 : 5 in PBS and was incubated at 37°C for 30 minutes with agitation. Each time, the mixture of 1 μg of rIL-2 and 5 μg of anti-mouse IL-2 in a volume of 150 μl was injected intraperitoneally (IP). Injection of IL-2C was started from the 3^rd^ week, daily in 3 consecutive days and then followed by once a week until the end of the experiment. A hyperbaric oxygen chamber for animal study was purchased from Particla (Daejeon, South Korea). The HBO protocol was conducted with 100% O_2_ at 3 ATA for 90 minutes after 20 minutes of compression, and then followed by 60 minutes of decompression. HBO treatment was given 5 times a week from the start to the end of the experiment. At the 14^th^ week of the experiment, intraperitoneal glucose tolerance test (IPGTT), and at the 15^th^ week, intraperitoneal insulin tolerance test (IPITT) were done. At the 16^th^ week, the mice were ethically sacrificed under general anesthesia, and spleen, liver and adipose tissues were dissected for histological and immunological study. For the study of WAT, epididymal white adipose tissues (eWAT) were dissected. For the BAT, interscapular brown adipose tissues (iBAT) were dissected. For the subcutaneous white adipose tissue, anterior subcutaneous adipose tissues were dissected from the upper left ventral region and inguinal subcutaneous adipose tissues were dissected.

### Glucose metabolism study

For the glucose tolerance tests (GTT), mice were intraperitoneally (IP) injected with 2 g/kg glucose (Sigma Aldrich, Saint Louis, MO, USA) after fasting for 15 hours. Blood glucose levels were measured with a glucometer (Accu-Check Performa kit, Roche, Basel, Switzerland) at 0, 15, 30, 60 and 120 minutes. For the insulin tolerance tests (ITT), mice were IP administered with 0.75 IU/kg insulin glargine (LantusTM, Sanofi, Paris, France) after fasting for 4 hours, and blood glucose levels were measured at 0, 15, 30, 60 and 120 minutes. The area under the curve (AUC) was calculated by PRISM program v5.01 (GraphPad Software Inc., La Jolla, CA, USA). Insulin levels in fasting sera were measured by using an insulin ELISA kit (ALPCO, Salem, NH, USA).

### Biochemical tests for lipid

Serum levels of total cholesterol (TC) and triglyceride (TG) were measured by using TC assay kit and serum TG quantification kit purchased from Cell Biolabs, Inc. (San Diego, CA, USA) according to the manufacturer’s instructions.

## Histology

### H&E, Masson’s trichrome, Oil red O staining and microscopic analysis

The dissected tissue was fixed in 10% formalin. Serial sections (4 μm) were mounted on slides and stained with hematoxylin and eosin (H&E). Adipocyte size and number of eWAT, as well as the thickness of anterior subcutaneous WAT were measured by Image J program (National Institutes of Health, Bethesda, MD, USA).

Masson’s trichrome staining was applied for the detection of collagen fibers in the liver tissues. Paraffin-embedded liver sections were stained with Weigert’s iron hematoxylin working solution (Polysciences inc., Warrington, PA, USA) for 10 minutes and then with Biebrich scarlet-acid fuchsin solution (Sigma Aldrich) for 10 minutes. After washed with D.W., the sections were differentiated in phosphomolybdic-phosphotungstic acid solution (Sigma Aldrich) for 10 minutes and were stained with aniline blue solution (Sigma Aldrich) for 5 minutes. Then, the tissue samples were dehydrated and mounted.

Oil red O staining was applied for the evaluation of fat accumulation in the liver. Freshly dissected liver tissues were embedded in Tissue-Tek O.C.T. compound (Thermo Fisher Scientific, MA, USA), and were frozen at -196.5°C. The frozen sections (10 μm thickness) were prepared and dried by air. The slides were rinsed with 60% isopropanol (Sigma Aldrich) and were stained with Oil red O solution (Sigma Aldrich) for 15 minutes. The slides were rinsed again with 60% isopropanol and were stained with alum hematoxylin (Polysciences inc.). The slides were rinsed with D.W. and were mounted with aqueous mount solution (Thermo Fisher Scientific).

For the assessment of NAFLD activity score (NAS), randomly selected 3 independent images were examined and scored for the steatosis, lobular inflammation, and ballooning. The NAS is calculated by the sum of scores for steatosis (0 - 3), lobular inflammation (0 - 3), and hepatocyte ballooning (0 - 2), and the total score ranges from 0 to 8 [42].

### Immunohistochemistry and quantification of expression levels

The paraffin-embedded tissue sections (4 μm) were hydrated in ethanol and D.W. Tissue slides were heated in boiling water to retrieve antigens for 10 - 20 minutes. Endogenous peroxidase was blocked by treatment with H_2_O_2_ containing peroxidase blocking reagent (DAKO, Santa Clara, CA, USA) for 10 minutes in the dark. After washed, the sections were treated with those primary antibodies such as F4/80 (Abcam, Cambridge, UK, 1:100 dilutions), HIF-1α (Bethyl Laboratories, Inc., Montgomery, TX, USA, 1:100 dilutions), UCP-1 (Abcam, 1:4,000 dilutions) for 1 - 2 hours at RT and then treated with HRP polymer (Cell Signaling Technology, Inc., MA, USA) for 30 minutes. Then, the sections were treated with DAB chromogen (Biocare Medical, Pacheco, CA, USA) for 5 minutes at RT in the dark and then stopped by D.W. The slides were counter-stained with hematoxylin for 5 sec. Tissue sample was dehydrated and mounted.

For the quantitative analysis of HIF-1α expression, the slides were applied for high-resolution whole slide scan using the Vectra Polaris (PerkinElmer Inc., Hopkinton, MA, USA). The regions for analysis were randomly selected 3 - 5 spots and applied to the inForm (Image Analysis Software, Perkin Elmer). Adipose and liver tissues were selected by tissue and blank segmentation algorithm. Among those tissue areas, cell segmentations were performed by nucleus intensity and cell morphology. The inForm generated intensity reports for DAB markers from the each analyzed cells. The DAB positive cells were measured in the selected 3 - 5 images per a group at the resolution of X10 in the eWAT and X20 in the liver.

### Flow cytometry

For the preparation of splenocytes, spleens were teased and minced, and were treated with ACK solution (Sigma Aldrich) for 5 minutes at RT, and then filtered through 70 μm cell strainers (SPL Life Sciences Co., Korea). For enrichment, the cells were spun on a 17.5% Accudenz (Accurate Chemical & Scientific, Westbury, NY, USA) density gradient centrifugation (2,000 rpm, 20 minutes, 20°C). Interface layer and pellet were used for the staining of macrophages and T cells, respectively. Stromal vascular fractions (SVFs) were prepared from eWAT by using adipose tissue dissociation kit (Miltenyi Biotec, Bergisch Gladbach, Germany) according to the manufacturer’s instruction. After the fractions were prepared as single cells, they were stained for M1 macrophages and Tregs using anti-F4/80 FITC (BM8, eBioscience) and anti-CD11c PE (HL3, BD, Franklin Lakes, NJ, USA) after Fc blocking with anti-CD16/32 antibodies (2.4G2, BD). The cells were also stained with anti-CD3 PerCP/Cy5.5 (17A2, Biolegend, San Diego, CA, USA), anti-CD4 FITC (GK1.5, Biolegend) and anti-CD25 Brilliant Violet 711 (PC61, Biolegend) antibodies. For the staining of intranuclear FoxP3, the cells were permeabilized with FoxP3 staining buffer solution (eBioscience) and then stained with anti-FoxP3 APC (FJK-16s, eBioscience). The stained cells were analyzed using LSRFortessa (BD) and FlowJo software v10 (Tree Star, Ashland, OR, USA).

### Statistical analysis

Results are presented as the mean ± SEM. Statistical significance was evaluated with *t*-test and two-way ANOVA. Significance was set at * *p*<0.05, ** *p*<0.01, *** *p*<0.001. The significance of each experimental group was evaluated by comparison with the control group.

## Results

### Attenuation of the body weight gain by the individual or combination treatment with IL-2C and HBO

Body weight gain was faster and more prominent in the HFD groups than in the LFD groups. Body weights in the HFD control group became heavier than in the LFD control group from the 4^th^ week and the difference between the groups became progressively more prominent until the end of the experiment (S1 Fig). At the end of the experiment, the body weights in the groups treated with the individual or combination of the IL-2C and HBO were significantly lower than those in the controls of the HFD group (Fig 1A). In the LFD groups, at the end of the experiment, the body weights in the group by individual treatment with IL-2C and combination treatment were also significantly lower than those in the controls. The body weights in the group by individual treatment with HBO was also lower but was not statistically significant. There was no significant difference in the food intake between the experimental groups (Fig 1B).

**Fig 1.**
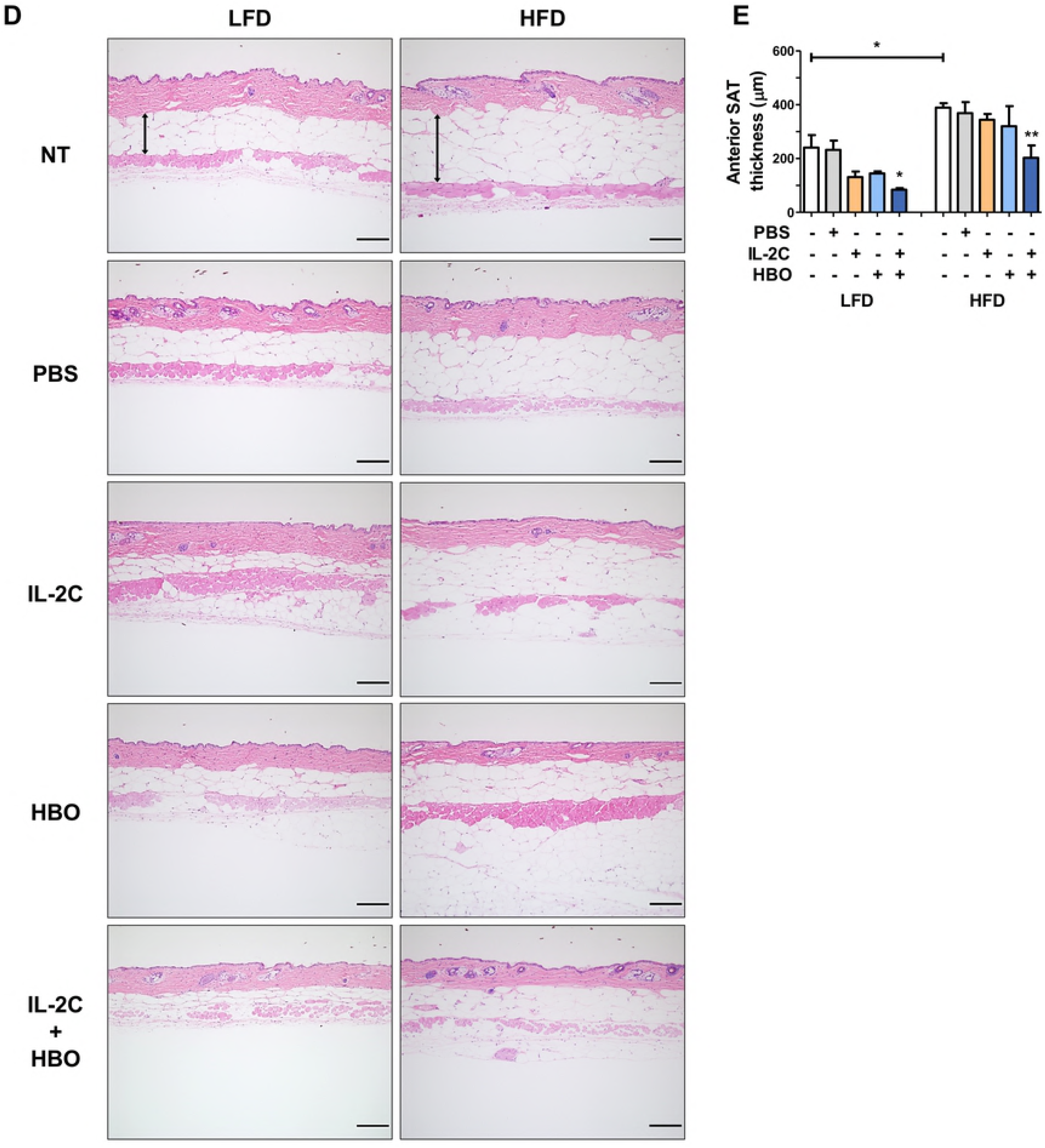
Body weights and food intake. Body weights of the experimental groups at the end of the experiment (A). Food intake (B).

### Improvement of impaired glucose metabolism and dyslipidemia by the individual or combination treatment with IL-2C and HBO

At the end of the experiment, blood glucose levels after fasting for 15 hours were significantly higher in the HFD control groups than in the LFD groups, suggesting hyperglycemia was induced (Fig 2A). Individual treatment with IL-2C or HBO decreased blood glucose levels, but without statistical significance, while combination treatment decreased it significantly. IPGTT showed delayed glucose clearance from the blood in the HFD control groups, suggesting dysregulation of glucose metabolism (Fig 2B). In the combination treatment groups, the blood glucose levels were significantly decreased starting from 30 minutes and from 60 minutes in each LFD and HFD compared to the control groups (Fig 2C and D). Glucose clearance rate after glucose injection was shortened by the individual or combination treatment with IL-2C and HBO (Fig 2E). In order to investigate if insulin functions appropriately, blood glucose levels were measured after IP injection of insulin. The blood glucose levels after fasting for 4 hours was significantly lower in the HFD groups by individual treatment with HBO or combination treatment (Fig 2F). The levels of blood glucose at 2 hours after insulin injection were showed significantly higher and delayed glucose clearance in the HFD control groups, therefore this result suggests the inappropriate function of insulin (Fig 2G). Individual or combination treatment with IL-2C and HBO decreased blood glucose levels after injection of insulin, especially combination treatment was significantly lower during 2 hours in the HFD groups, but there was no significant change in the LFD groups (Fig 2H and I). Therefore, glucose clearance rate after insulin injection was significantly improved by the individual or combination treatment with IL-2C and HBO in the HFD (Fig 2J). In addition, serum insulin levels after fasting for 15 hours were significantly higher in the HFD control groups (Fig 2K). *Albeit* at higher insulin levels, hyperglycemia was not corrected, suggesting insulin resistance was induced in the HFD control groups. Individual or combination treatment with IL-2C and HBO not only decreased fasting serum insulin levels in the HFD groups, in parallel with the decrease of fasting blood glucose levels but also enhanced glucose clearance from the blood after injection of insulin, suggesting improvement of insulin function. At the end of the experiment, hypercholesterolemia was induced in the HFD control groups, which was improved by the individual or combination treatment with IL-2C and HBO (Fig 2L). Meanwhile, serum triglyceride levels were not significantly different between the LFD and HFD groups (Fig 2M). Taken together, it was suggested that the individual or combination treatment with IL-2C and HBO improved impaired glucose metabolism and dyslipidemia induced by HFD.

**Fig 2.**
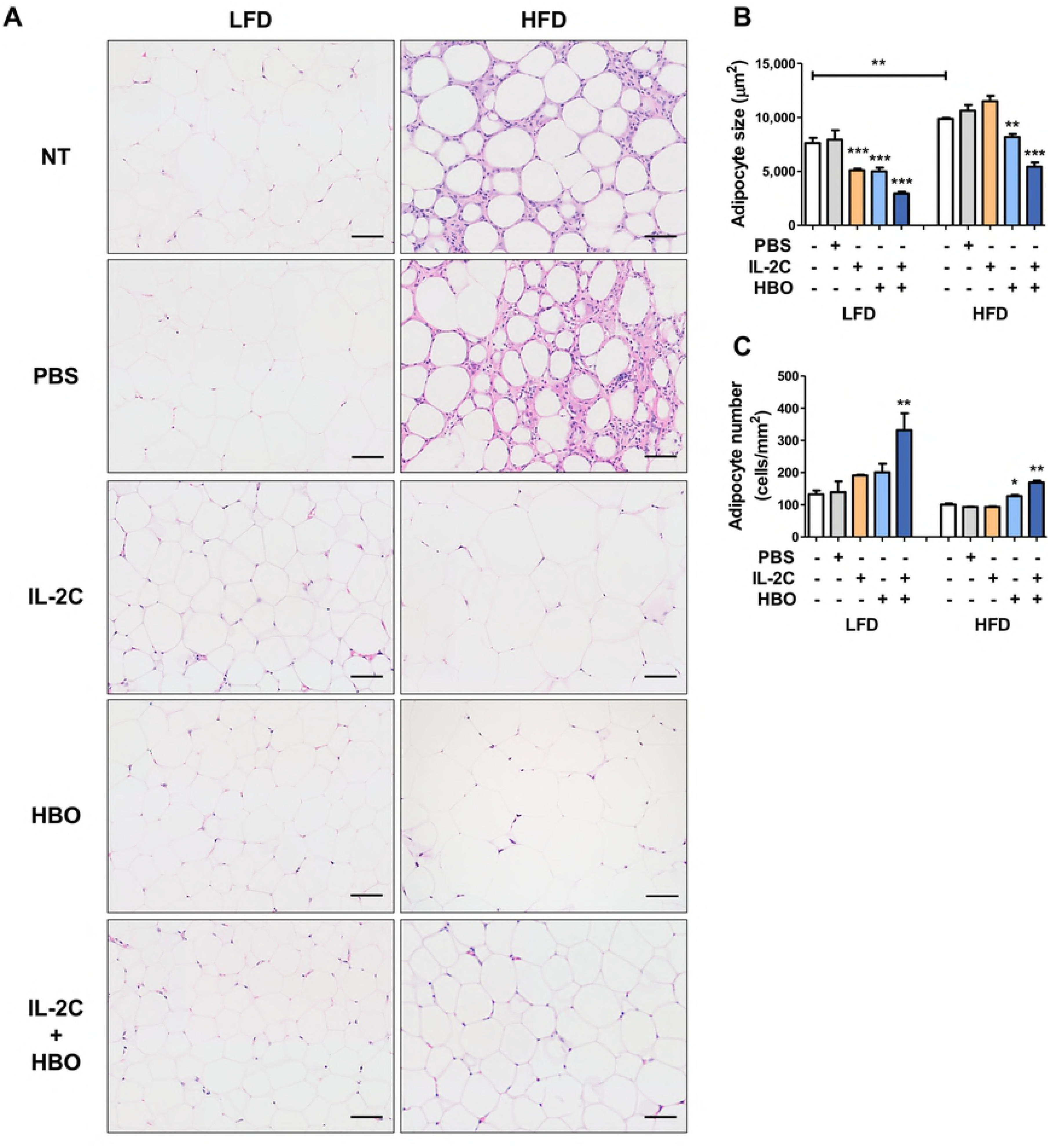
Blood and biochemical tests to analysis of glucose and lipid metabolism. IPGTT and IPITT were conducted in the experimental groups of LFD and HFD. Blood glucose levels at the start (after fasting for 15 hours) (A) and at the end (2 hours after glucose injection) (B) of IPGTT. The levels of blood glucose in the LFD (C) and HFD groups (D) during the 2 hours, and AUC of IPGTT (E). Blood glucose levels at the start (after fasting for 4 hours) (F) and at the end (2 hours after the insulin injection) (G) of IPITT. The levels of blood glucose in the LFD (H) and HFD groups (I) during the 2 hours, and AUC of IPITT (J). The levels of insulin (K), total cholesterol (L) and triglyceride (M) in the sera after fasting for 15 hours.

### Modulation of HFD-induced adipose tissue remodeling by the individual or combination treatment with IL-2C and HBO

As obesity progresses by HFD, adipose tissues undergo dynamic remodeling by changing the size and number of adipocytes. Histological examination of the eWAT showed many enlarged adipocytes in the HFD control groups, but many small or shrunken cells were also observed (Fig 3A). Around the small adipocytes, many inflammatory cells were infiltrated, showing many typical crown-like structures. The groups by the individual or combination treatment with IL-2C and HBO has scarcely observed the infiltration of inflammatory cells, and the size of adipocytes was rather homogenous with less variation in the eWAT. The size of adipocytes in the HFD control groups was larger than LFD groups (Fig 3B). It was decreased by the individual or combination treatment in both LFD and HFD groups. On the other hand, the size of the adipocytes in the HFD groups by individual treatment with IL-2C was not different from that in the control groups. Inversely proportional to size, the number of adipocytes in the HFD control groups decreased in the same area compared with the treatment groups (Fig 3C). For the evaluation of fat accumulation in the subcutaneous adipose tissue (SAT), anterior SAT thickness was measured from the ends of the dermis to muscle layer (Fig 3D and E). The depth of anterior SAT was also significantly thicker in the HFD control groups compared with the LFD groups. Meanwhile, it was significantly thinner in the LFD and HFD groups by combination treatment with IL-2C and HBO compared with that in the control groups. Therefore, HFD-induced adipose tissue remodeling was prevented in both VAT and SAT by the individual or combination treatment with IL-2C and HBO.

**Fig 3.**
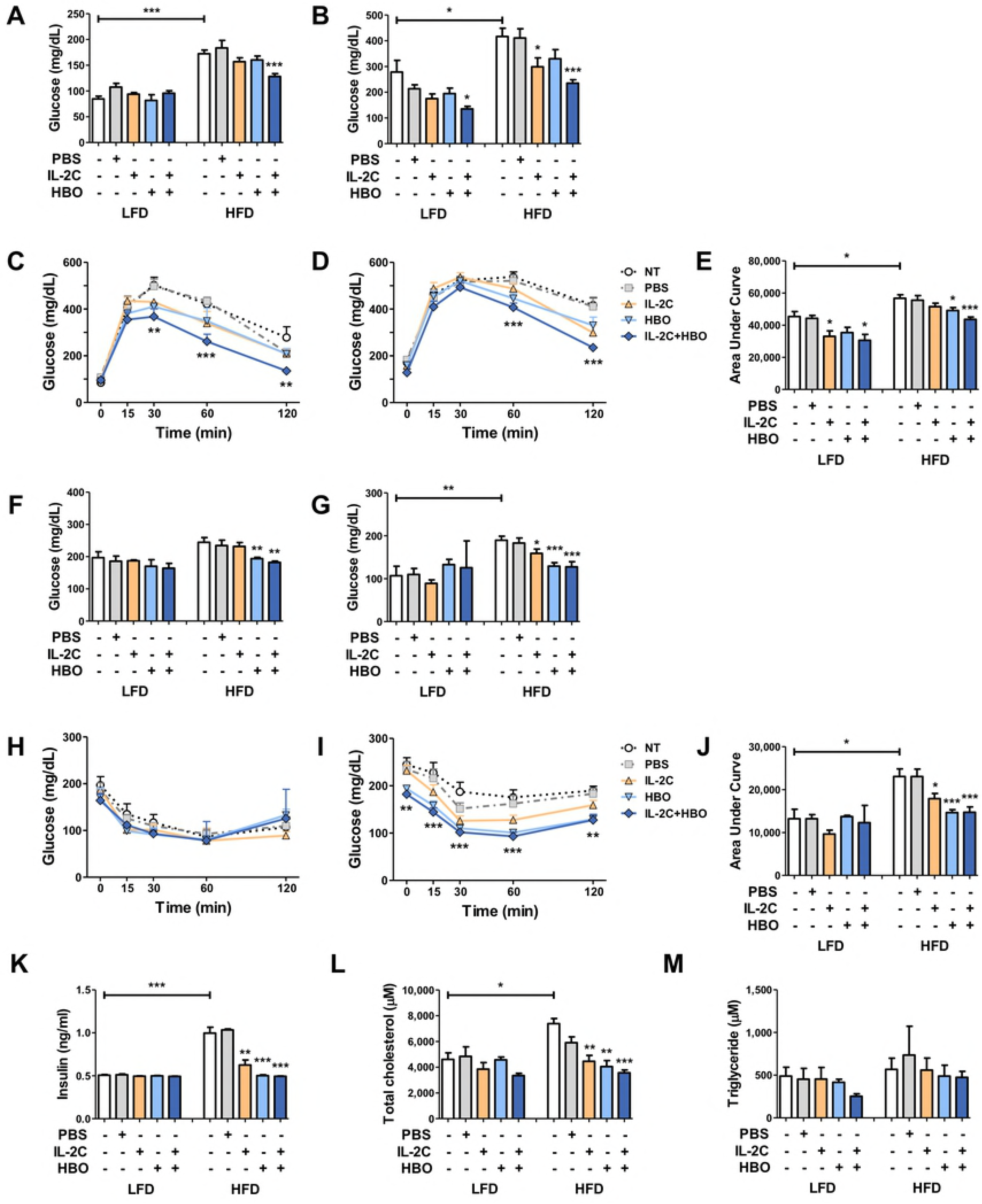
Histological analysis of HFD-induced adipose tissue remodeling in the VAT and SAT. H&E staining of eWAT (X200, scale bar is 100 μm) shows hypertrophic adipocytes and crown-like structures in the HFD control groups (A). The groups by the individual or combination treatment with IL-2C and HBO has reduced the size of adipocytes, and immune cells infiltration in the eWAT. Histological analysis of the size (B) and number (C) of adipocytes was performed by ImageJ program. H&E staining shows the side of longitudinally sectioned anterior SAT (X100, scale bar is 200 μm) (D). ‘Bidirectional arrows’ indicate adipose thickness from the ends of the dermis to muscle layer. The thickness of anterior SAT was analyzed using the ImageJ program by measuring the length of arrows mentioned above (E).

### Improvement of non-alcoholic fatty liver disease (NAFLD) by the individual or combination treatment with IL-2C and HBO

The liver plays important roles in the lipid metabolism such as import of serum FFAs and generation, store and then export of lipids and lipoproteins. In addition, it was reported that 80% of obese people have the NAFLD. Histological examination of the H&E stained liver sections in the HFD control groups showed severe steatosis with many microvesicular and macrovesicular fatty changes (Fig 4A). Oil red O staining showed more evident fat accumulation in the liver of the HFD control groups, but it was dramatically reduced by the individual or combination treatment with IL-2C and HBO (Fig 4B). According to the fat accumulation in the liver, the weight of liver was significantly increased by HFD, which were also significantly decreased by individual treatment with IL-2C and combination treatment, but not significantly with individual treatment with HBO (Fig 4C). There was no significant difference between in the LFD each group. In the HFD control groups, hepatocyte ballooning and Mallory-Denk bodies were also observed, suggesting degenerative changes of the liver (Fig 4D). Lobular inflammation by infiltration of polymorphonuclear leukocytes and lymphocytes also suggested ongoing hepatitis. Masson’s trichrome staining shows slight but evident fibrotic changes (Fig 4E). Taken together, it could be argued that non-alcoholic steatohepatitis (NASH) has been induced by HFD, and semi-quantitative analysis also showed the significant increase of NAFLD activity score (NAS) in the HFD control groups (Fig 4F-I). Meanwhile, the histological characteristics of NASH were dramatically attenuated in parallel with the significant decrease of NAS in the groups by the individual or combination treatment with IL-2C and HBO.

**Fig 4.**
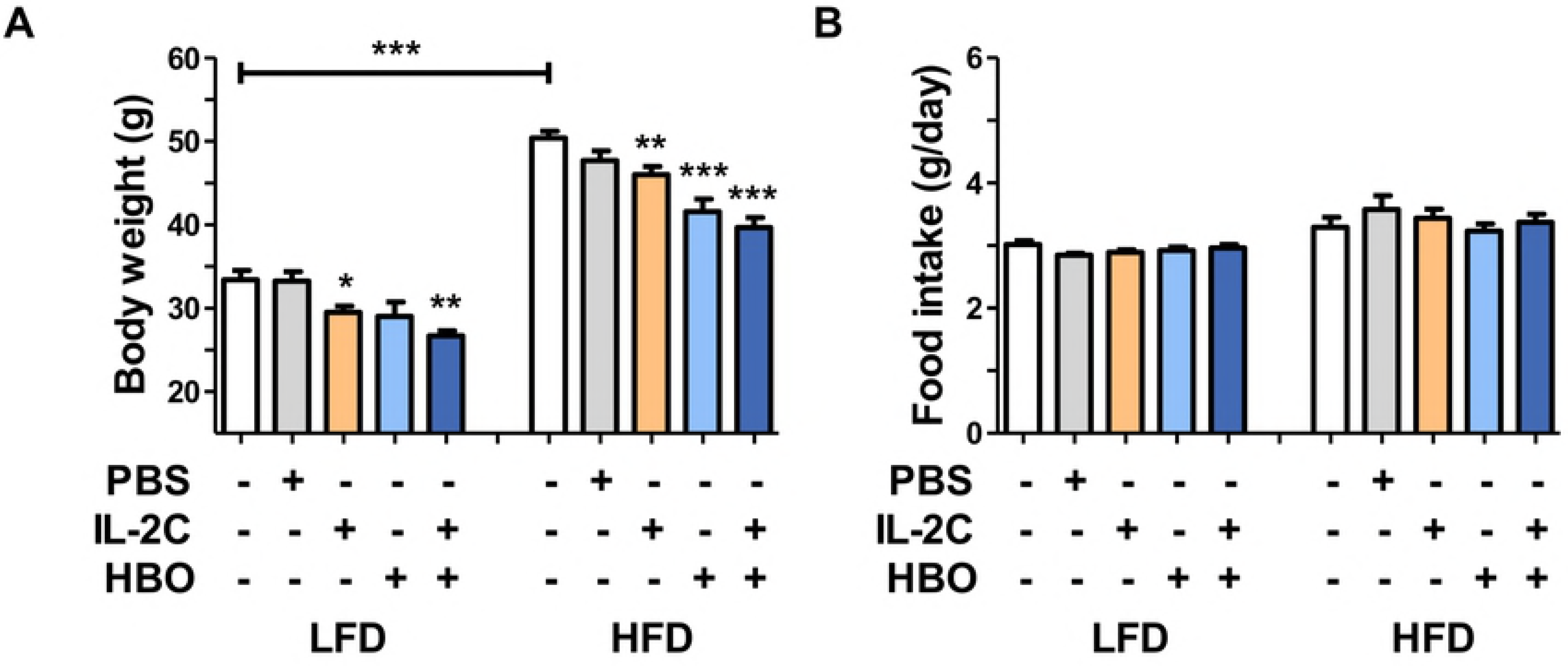
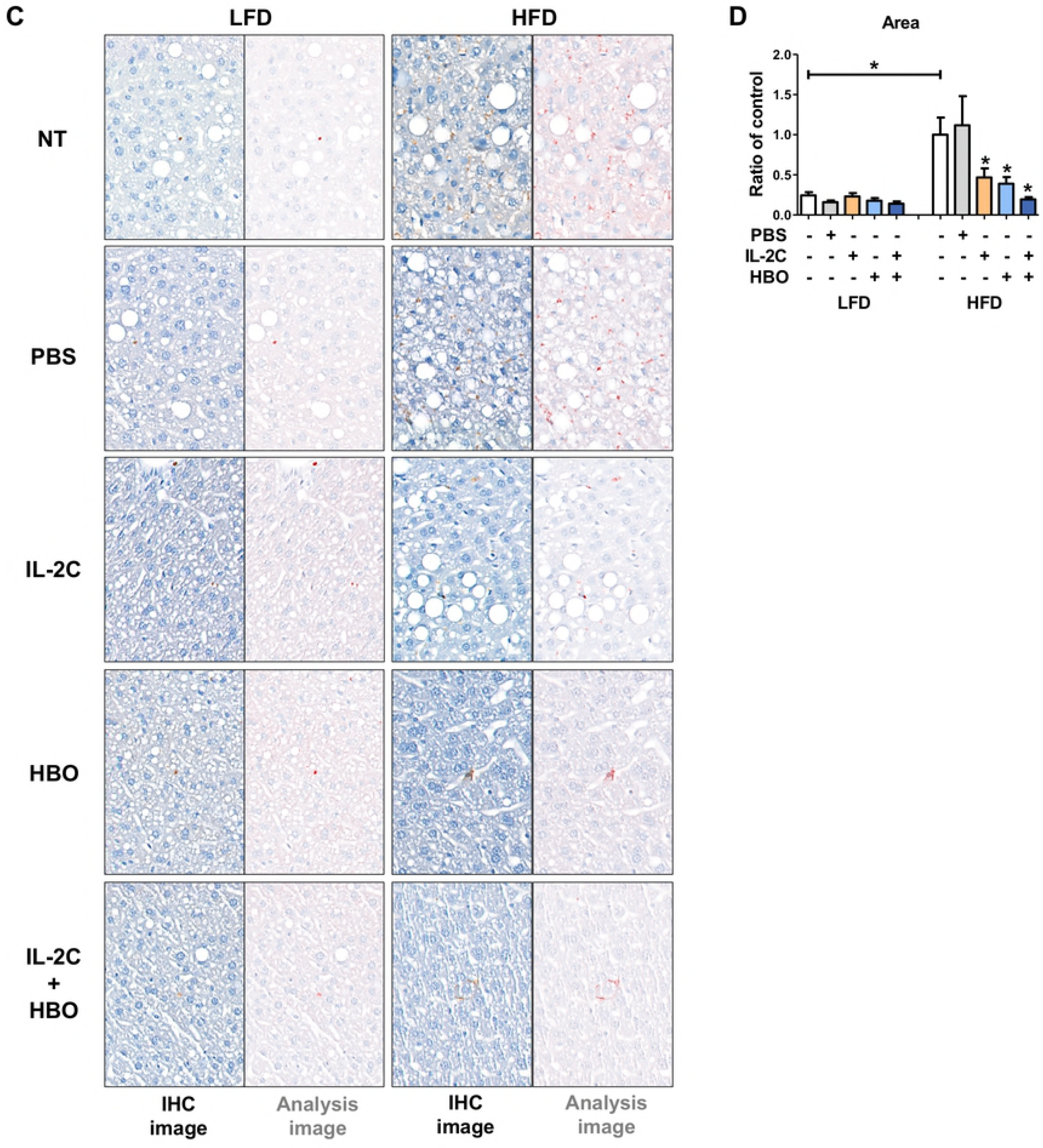
Histopathological analysis of NAFLD induced by HFD. H&E staining of the liver in the experimental groups (X200, scale bar is 100 μm) (A). H&E staining shows severe steatohepatitis in the HFD control groups, but it was prevented in the groups by the individual or combination treatment with IL-2C and HBO. Oil red O staining (X200, scale bar is 100 μm) (B) shows the fat depositions in the liver. Fat vacuoles stained red were overall scattered in the liver of HFD control groups, but the fat vacuoles by the individual or combination treatment with IL-2C and HBO become small and less. The weight of the liver (C). H&E staining of the HFD control groups (X400) (D). ‘Blue arrows’ indicate the Mallory-Denk bodies, ‘* and **’ indicate the microvesicular and macrovesicular fatty changes, and ‘Yellow arrow’ indicates lobular inflammation composed of PMLs and lymphocytes. Masson’s trichrome staining shows fibrotic changes in the HFD control groups (X400) (E). Semi-quantitative analysis was performed to assess of steatosis (F), lobular inflammation (G), and hepatocyte ballooning (H) for NAFLD activity score (NAS) (I).

### Suppression of HFD-induced inflammation by the individual or combination treatment with IL-2C and HBO

IHC and flow cytometry was conducted to evaluate the inflammatory state induced by obesity. Firstly, the IHC study for the F4/80 was conducted with the analysis of the infiltrated macrophages in the eWAT. As a result, many macrophages composing the crown-like structures were infiltrated in the eWAT of the HFD control groups, but it was less in the mice treated with the individual or combination of IL-2C and HBO (Fig 5A and B). Flow cytometry was performed to evaluate the inflammatory state of adipose tissue and spleen, and the proportional changes of M1 macrophages and Tregs were examined locally and systemically. Flow cytometric analysis also showed the significant increase in F4/80^+^CD11c^+^ M1 macrophage proportion among the VSFs of eWAT in the HFD control groups (Fig 5C). Meanwhile, the proportion of M1 macrophage was significantly decreased by combination treatment with IL-2C and HBO. Individual treatment of IL-2C or HBO in the HFD groups were also decreased but without significance. Flow cytometric analysis of the splenocytes also showed the similar pattern of increase the M1 macrophage proportion in the HFD control groups and decrease by the individual or combination treatment with IL-2C and HBO, and the strength of statistical significance was stronger than eWAT (Fig 5D). In the eWAT as well as in the spleen, combination treatment with IL-2C and HBO in the HFD groups significantly reduced the proportion of M1 macrophage. The CD4^+^FoxP3^+^ Treg proportion in both eWAT and spleen of the HFD groups was significantly increased by the individual or combination treatment with IL-2C and HBO. The proportion of Treg among the VSFs of eWAT was significantly decreased in the HFD control groups compared with that in the LFD groups (Fig 5E). In contrast, the proportion of Treg in the HFD groups was increased by the individual or combination treatment with IL-2C and HBO, and the proportion of Treg in the LFD groups increased only in the individual treatment with HBO. The proportion of Treg in the spleen of the HFD control groups was also decreased compared with the LFD groups (Fig 5F). It was significantly increased in both LFD and HFD groups by individual treatment with IL-2C and combination treatment. In summary, in both eWAT and spleen of the HFD control groups, M1 macrophages were increased whereas Tregs were decreased compared with the LFD control groups, suggesting the pro-inflammatory state. On the other hand, the individual or combination treatment with IL-2C and HBO reversely changed the proportion of the immune cells induced by HFD, suggesting suppression of the inflammatory responses.

**Fig 5.**
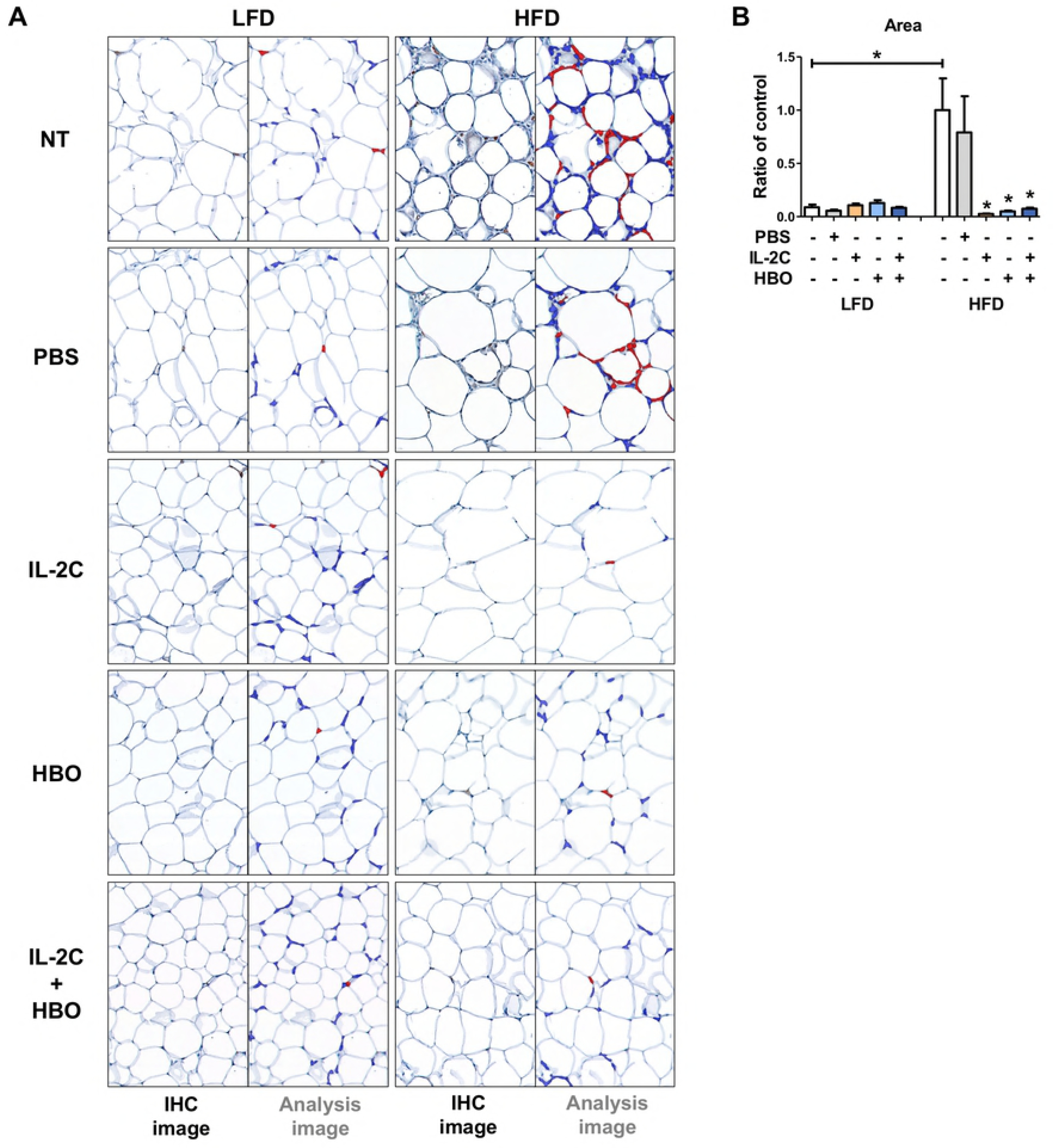
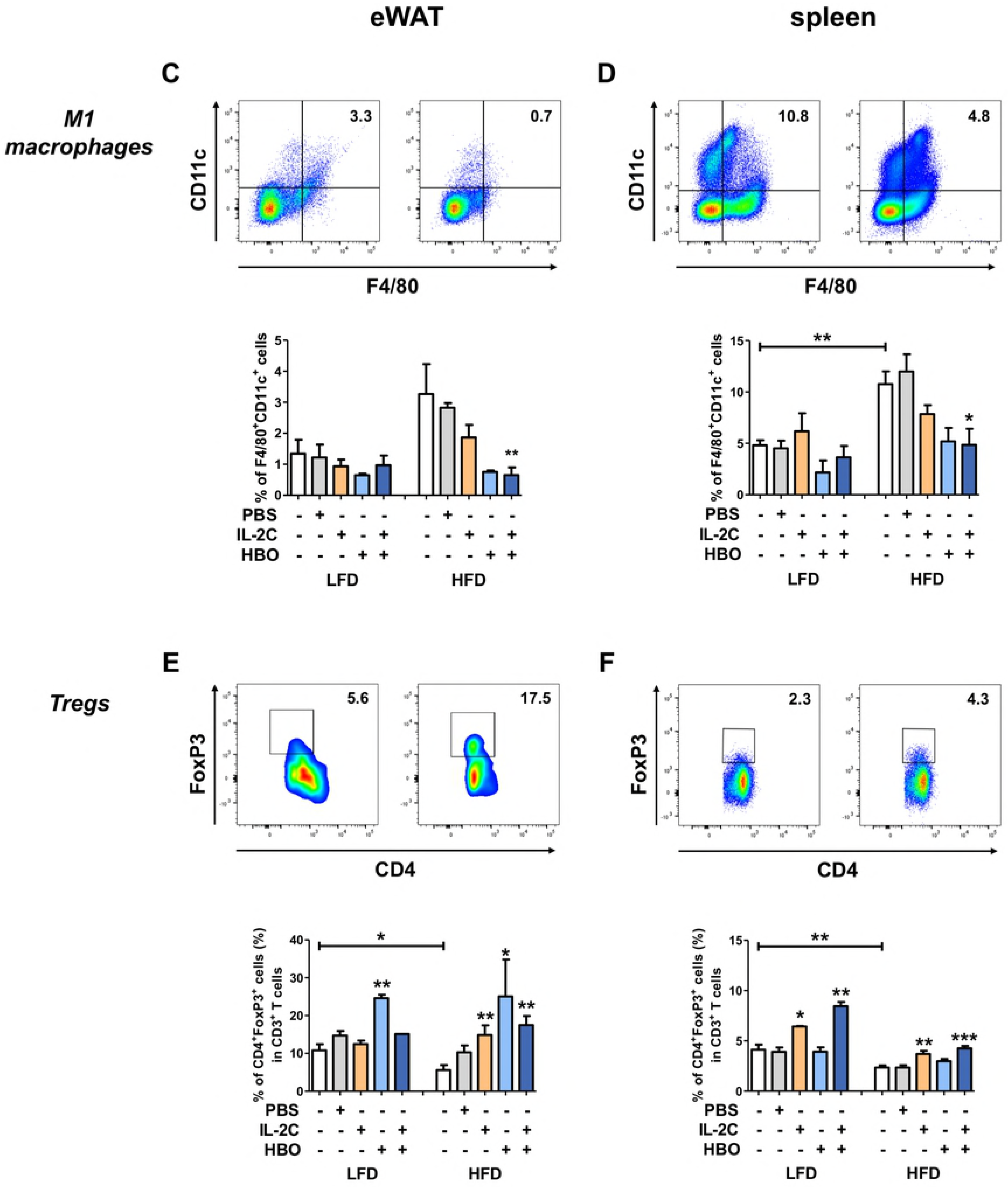
Histopathology and immunological analysis of eWAT and spleen induced by HFD. IHC for F4/80 as a macrophages marker in the eWAT (X400, scale bar is 50 μm) (A). Quantification of F4/80 positive macrophage cells count (B). Flow cytometry for M1 macrophages and Tregs population in the eWAT and spleen. F4/80 and CD11c double positive cells were gated as M1 macrophages in the eWAT (C), and spleen (D). CD4 and FoxP3 double positive cells in CD3^+^ T cells were gated as Tregs in the eWAT (E), and spleen (F).

### Reduction of HIF-1α expression in the eWAT and liver by the individual or combination treatment with IL-2C and HBO

Another cardinal manifestation of obesity is hypoxia in the WAT. IHC showed scarce expression of HIF-1α in the eWAT of the LFD groups, which was significantly enhanced in the HFD control groups, suggesting hypoxic state (Fig 6A and B). The enhanced expression of HIF-1α in the HFD control groups was significantly decreased to the comparable levels with that in the LFD groups by the individual or combination treatment with IL-2C and HBO, suggesting the normoxic state. IHC for HIF-1α in the liver tissues showed a similar pattern of increase of HIF-1α expression in the HFD control groups, which was decreased by the individual or combination treatment with IL-2C and HBO (Fig 6C and D).

**Fig 6.**
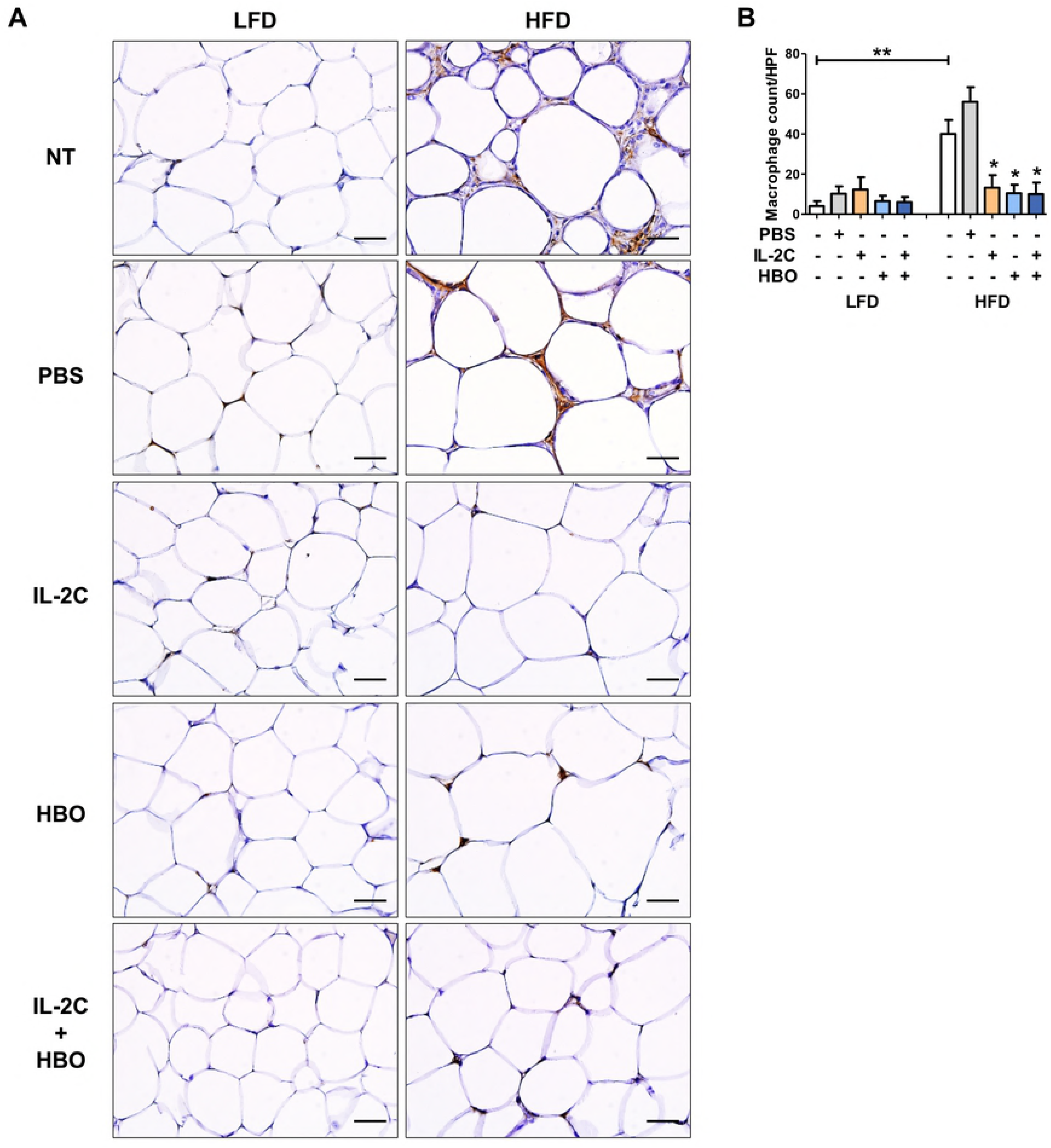
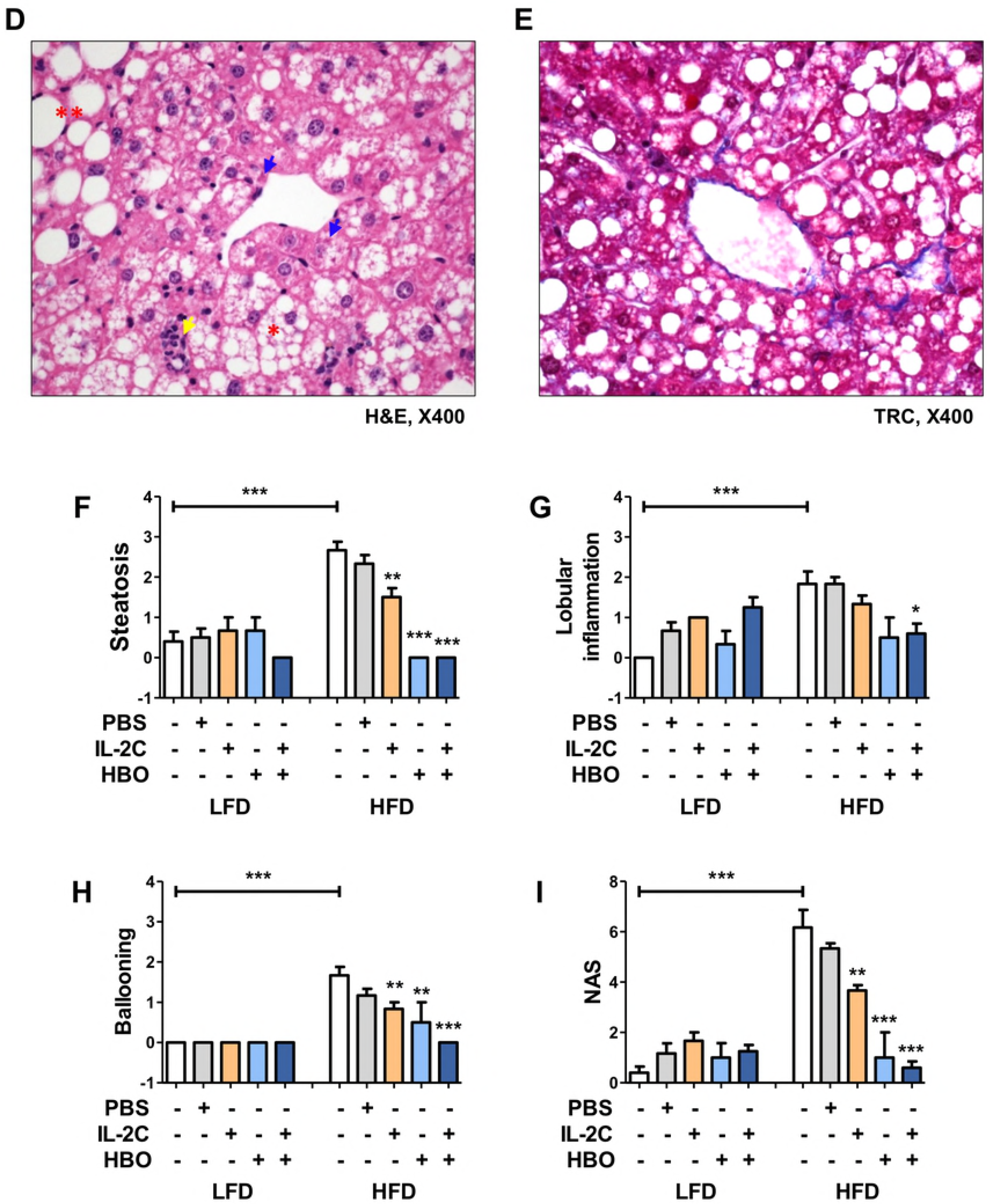
Immunohistochemistry and analysis for HIF-1α in the eWAT and liver. Immunohistochemistry for HIF-1α in the eWAT and quantification of HIF-1α expression area. The left of the picture is the IHC staining image and the right paired picture is the analysis image (A). Quantification the ratio for HIF-1α positive area compared with HFD control groups (B). Immunohistochemistry for HIF-1α in the liver and quantification of HIF-1α expression area. (C). Quantification the ratio for HIF-1α positive area compared with HFD control groups (D).

### Activation of adipose tissue browning by the individual or combination treatment with IL-2C and HBO

Histological examination of the iBAT showed small and multilocular features in the LFD groups (Fig 7A). In the HFD control groups, many unilocular hypertrophic adipocytes were observed with infiltration of inflammatory cells, suggesting whitening of adipose tissue. These histological features were less prominent by the individual or combination treatment with IL-2C and HBO of the HFD. The weight of iBAT was also slightly increased in the HFD control groups, compared with the LFD control groups, but without statistical significance (Fig 7B). In the mice treated with the combination of IL-2C and HBO, the weight of iBAT was significantly decreased in both LFD and HFD groups compared with each control groups. IHC showed that the uncoupling protein-1 (UCP-1) was intensely expressed in the iBAT of the LFD groups, which became more intense by the individual or combination treatment with IL-2C and HBO (Fig 7C and D). The UCP-1 expression in the iBAT of the HFD control groups was slightly less intense, which was also strengthened by the individual or combination treatment with IL-2C and HBO.

In addition, histological examination of the iWAT of the LFD groups showed homogeneous unilocular adipocytes, it was almost similar to those of eWAT (Fig 7E). However, many small multilocular adipocytes were observed by the individual or combination treatment with IL-2C and HBO. In the HFD control groups, iWAT showed many hypertrophic adipocytes along with the infiltration of inflammatory cells. The size of iWAT of the HFD by the individual or combination treatment with IL-2C and HBO were smaller than the HFD control groups. The weight of iWAT was also decreased by combination treatment with IL-2C and HBO in the LFD groups (Fig 7F). In the HFD control groups, the weight of iWAT was remarkably increased compared with the LFD groups. It was restored to the comparable level with LFD by combination treatment with IL-2C and HBO. In addition, the expression of UCP-1 in the iWAT was scarcely expressed in the LFD control groups, but it was gradually increased by the individual or combination treatment with IL-2C and HBO (Fig 7G and H). It was higher by individual treatment with HBO in comparison to the individual treatment with IL-2C, and the combination treatment with IL-2C and HBO showed the highest increase in UCP-1 expression. UCP-1 expression was hardly observed in the HFD control groups, but it was only expressed in the group treated with combination of IL-2C and HBO. As a result, the whitening of the adipose tissue induced by the HFD was prevented by the individual or combination treatment with IL-2C and HBO, and it was activated the conversion to the browning of the adipose tissue.

**Fig 7.**
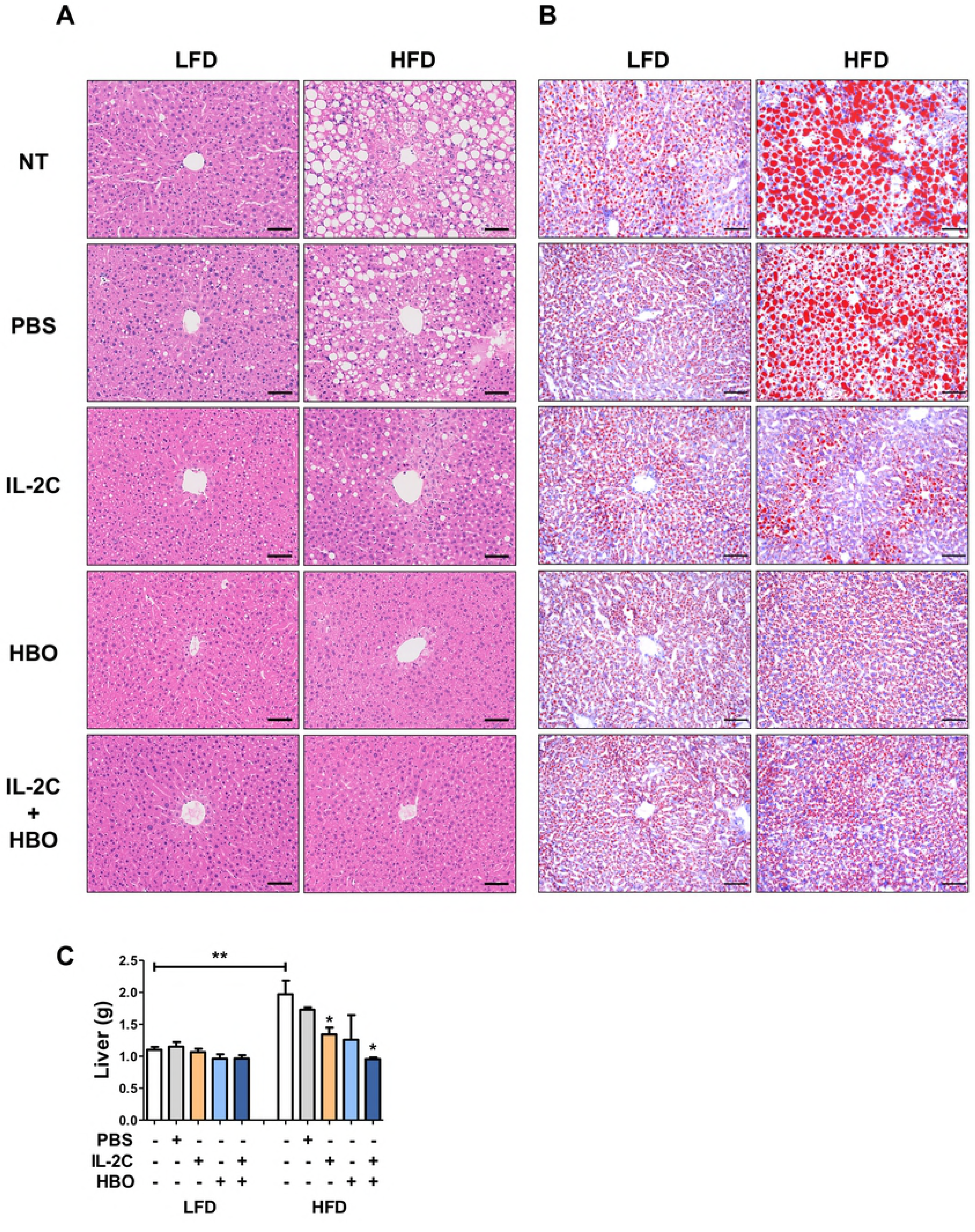
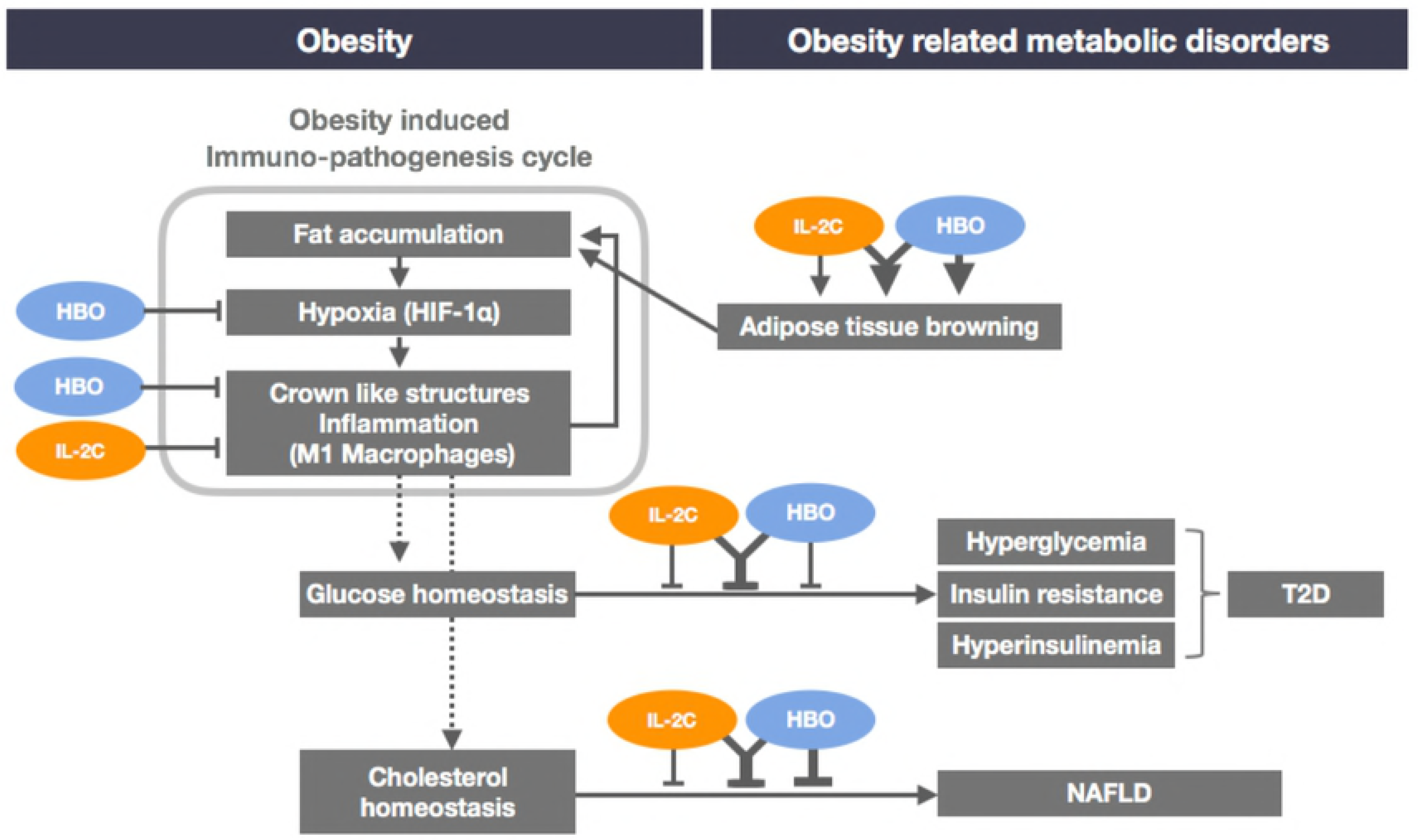
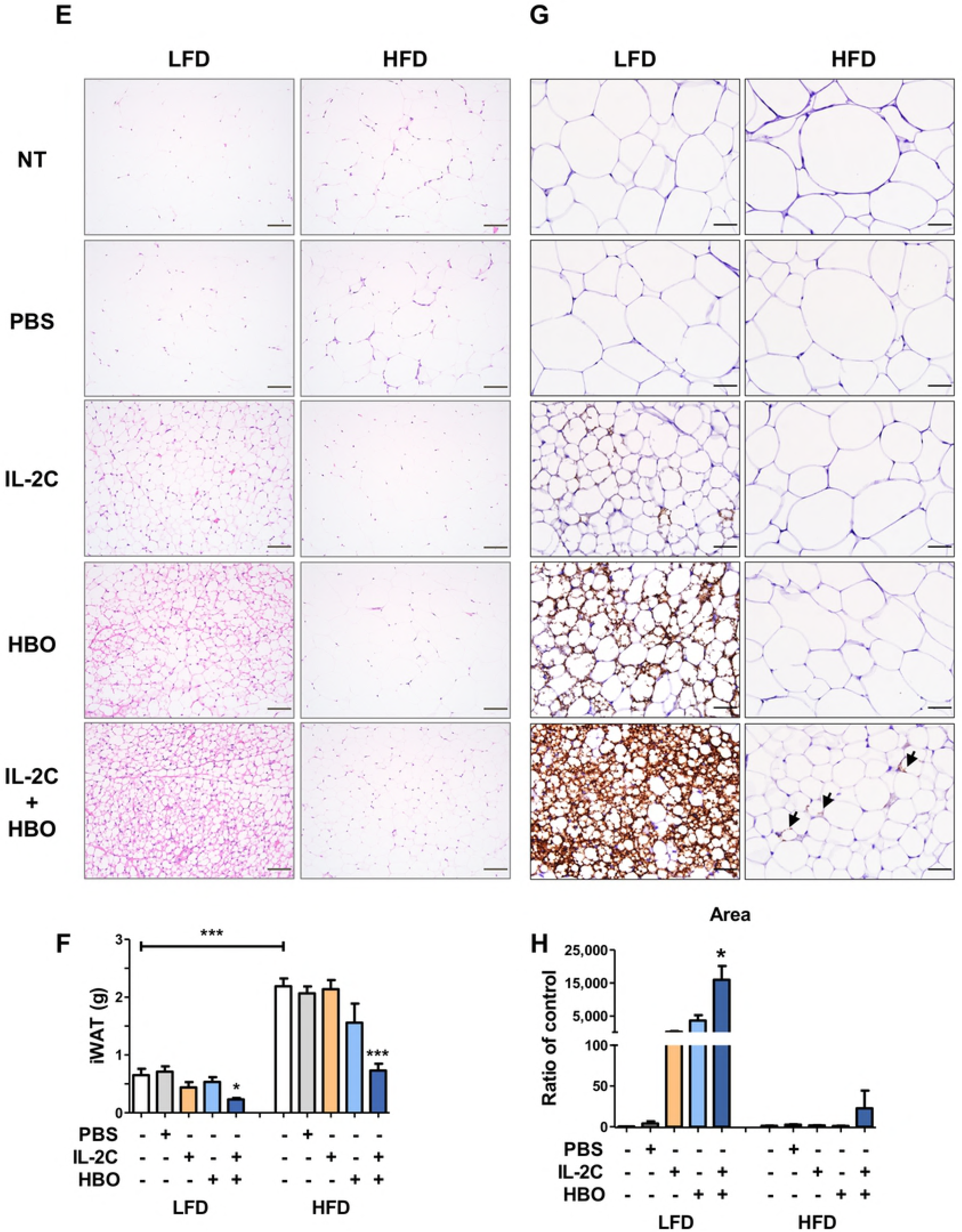
H&E staining and Immunohistochemistry for observation of adipose tissue browning in the iBAT and iWAT. H&E staining of the iBAT (X200, scale bar is 100 μm) shows adipose tissue whitening by the histomorphologic change from the multilocular to unilocular and increase of immune cells infiltration in the HFD control groups (A). Inversely, the individual or combination treatment with IL-2C and HBO are less fat accumulation and immune cells infiltration in the iBAT. The weight of the iBAT (B). IHC for UCP-1 in the iBAT (X400, scale bar is 50 μm) (C). Quantification the ratio for UCP-1 positive area compared with HFD control groups (D). H&E staining of the iWAT (X200, scale bar is 100 μm) shows hypertrophic adipocytes and immune cells infiltration in the HFD control groups, but it was reduced in the groups by the individual or combination treatment with IL-2C and HBO (E). In the LFD groups, adipose tissue browning was induced by individual treatment with IL-2C or HBO, and the group by combination treatment with IL-2C and HBO strongly induced browning. The weight of the iWAT (F). IHC for UCP-1 in the iWAT (X400, scale bar is μm) (G). ‘Arrows’ indicate the UCP-1 expression. Quantification the ratio for UCP-1 positive area compared with HFD control groups (H).

### Discussion

In the present study, the individual or combination treatment with IL-2C and HBO attenuated HFD-induced obesity and related metabolic disorders. The significant decrease of body weight gain by combination treatment with IL-2C and HBO was observed from as early as the 4^th^ week in the HFD group, and from the 8^th^ week in the LFD group. At the end of the experiment, decrease of body weight gain was as much as 21.2% and 20.1% in the HFD and LFD groups, respectively. Individual treatment with IL-2C or HBO also reduced weight gain, but the significant difference was always represented by combination treatment with IL-2C and HBO (S1 Table).

Impaired glucose metabolism in the HFD control groups was also improved by combination treatment with IL-2C and HBO. In the IPGTT, combination treatment with IL-2C and HBO not only significantly reduced the blood glucose levels both fasting, and 2 hours after glucose injection but also showed significantly decreased the AUC. In IPITT, blood glucose levels at 2 hours after insulin injection, AUC and fasting serum insulin levels were significantly decreased by individual treatment with IL-2C or HBO as well as by combination treatment. However, blood glucose levels after fasting for 4 hours were significantly decreased by individual treatment with HBO or combination treatment, but not by individual treatment with IL-2C. All the data presented in this study including body weight, blood glucose, serum insulin levels, and AUCs of IPGTT or IPITT, did not completely support the synergic effect between individual and combination treatment with IL-2C and HBO. However, combination treatment with IL-2C and HBO always represented significance.

Dyslipidemia was induced by HFD, which was also improved by the individual or combination treatment with IL-2C and HBO. Serum levels of total cholesterol were elevated by HFD, which was reduced by the individual or combination treatment with IL-2C and HBO. The size of the liver was remarkably increased to almost double and histological examination showed findings compatible with severe steatohepatitis in the HFD control groups. The combination treatment with IL-2C and HBO decreased the size of liver and improved steatohepatitis almost perfectly, in parallel with serum levels of total cholesterol. Considering that no pharmacological treatment has been approved so far for NASH, the novel findings in the present study might lead to the development of an innovative therapeutic strategy using the individual or combination treatment with IL-2C and HBO.

One of the underlying mechanisms by the individual or combination treatment with IL-2C and HBO for the altered glucose metabolism, dyslipidemia, and NASH might be related with the suppression of inflammation, while obesity is known as a systemic low-grade inflammatory state promoting the insulin resistance. F4/80^+^CD11c^+^ M1 macrophages known to dysregulate adipocyte signaling were increased in the HFD control groups locally in the adipose tissues as well as systemically in the spleen. By contrast, CD4^+^FoxP3^+^ Tregs that inhibits inflammation were decreased. Both IL-2C and HBO have been known to expand Tregs *in vivo*, in the present study, Tregs were increased by the individual or combination treatment with IL-2C and HBO locally in the adipose tissues as well as systemically in the spleen. In accordance with the increase of Tregs, M1 macrophages were decreased locally as well as systemically.

Another mechanism for the treatment effect might be related to the restoration of hypoxia. As adipocytes become hypertrophic, it also becomes short of blood supply and hypoxia. HIF-1α is a master mediator of hypoxic signal and is activated in obese adipose tissue. It is involved in the insulin resistance and fibrosis in the WAT [43]. Previously, it was reported that HIF-1α in myeloid cells promoted adipose tissue remodeling toward insulin resistance [44], but selective inhibition of HIF-1α ameliorated adipose tissue dysfunction [45]. In the liver, HIF-1α stimulates triglyceride accumulation by stimulation of lipin 1 expression [46]. As HIF-1α is closely linked to the glucose and lipid metabolism in both adipose tissue and liver, and therefore hyperoxia is one of the potential therapies for the treatment of hypoxia in obesity and diabetes [47]. In this study, individual treatment with HBO and combination treatment reduced HIF-1α expression in WAT and liver in the HFD groups, suggesting the hypoxic state is alleviated. In addition, individual treatment with IL-2C also reduced HIF-1α expression. it is not sure that how IL-2C is involved in the increase of oxygen tension in adipose tissue, but it can be speculated that weight reduction, as well as suppression of inflammation induced by treatment with IL-2C, may be related with the downregulation of HIF-1α.

Adipose tissue undergoes dynamic remodeling such as adipocyte hypertrophy and hyperplasia, alteration of adipokine secretion, and changes in the adipose tissue-resident cells in response to the change of the nutritional status. In this study, adipocyte hypertrophy and crown-like structures were evidently shown in the HFD control groups, and it was reduced by treatment with IL-2C and HBO. It was expected that the size of adipocyte was significantly reduced by the individual or combination treatment with IL-2C and HBO, in parallel with the decrease in body weight. However, the pattern of change of the visceral adipocyte size in the individual treatment with IL-2C was different from the expectation (Fig 3B). Nonetheless, glucose metabolism and insulin resistance were significantly improved by individual treatment with IL-2C. Furthermore, the thickness of SAT was significantly increased by HFD, and it was decreased by combination treatment with IL-2C and HBO. It was traditionally considered that VAT was the important organ for the development of insulin resistance. However, there are also studies reporting that SAT significantly influences the development of insulin resistance [48]. As SAT was increased by HFD, which was decreased by combination treatment with IL-2C and HBO, it could be considered that SAT remodeling may play an important role in the regulation of glucose and lipid metabolism.

In addition, fat browning is an important therapeutic target for non-shivering thermogenesis consuming calories. There are many studies about the relationship between BAT and glucose metabolism [49], and BAT also improves glucose homeostasis and insulin sensitivity in humans [50]. UCP-1 is an important molecule involved in the metabolic thermogenesis, and it is mainly expressed in the brown and beige adipose tissue. Interestingly, the individual or combination treatment with IL-2C and HBO in the HFD as well as LFD groups increased the UCP-1 expression at both the iBAT and iWAT. It is able to be a novel finding that the individual or combination treatment with IL-2C and HBO could be the alternative therapeutic strategy for the stimulation of fat browning.

In conclusion, the individual or combination treatment with IL-2C and HBO attenuated HFD-induced obesity and related metabolic disorders through suppression of inflammation and stimulation of fat browning. Therefore, the individual or combination treatment with IL-2C and HBO may be considered for the alternative therapeutic strategy for obesity and related metabolic disorders (Fig 8).

**Fig 8.**
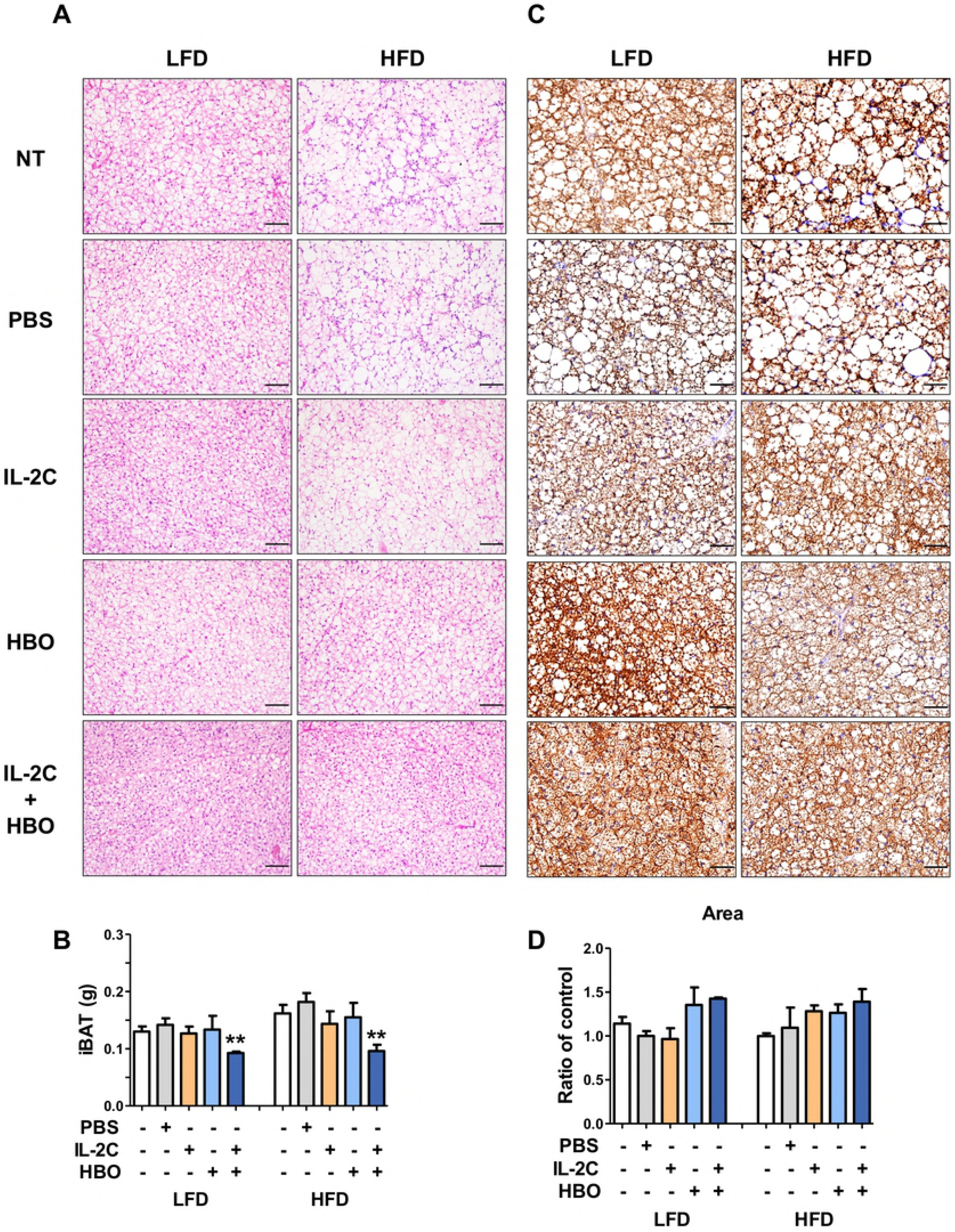
Mode of action. The individual and combination treatment of IL-2C and HBO in the prevention of HFD-induced obesity and related metabolic disorders.

## Supporting information

S1 Fig. The change of body weight in LFD (A) and HFD (B) groups during 14 weeks.

S1 Table. Summary of the individual and synergic effects of IL-2C and HBO in the HFD-induced obesity and related metabolic disorders.

## References

1. Romieu I, Dossus L, Barquera S, Blottiere HM, Franks PW, Gunter M, et al. Energy balance and obesity: what are the main drivers? Cancer Causes Control. 2017; 28(3):247–58. doi:10.1007/s10552-017-0869-z.

2. WHO, http://www.who.int/news-room/fact-sheets/detail/obesity-and-overweight.

3. Sahoo K, Sahoo B, Choudhury AK, Sofi NY, Kumar R, Bhadoria AS. Childhood obesity: causes and consequences. J Family Med Prim Care. 2015; 4(2):187–92. doi:10.4103/2249-4863.154628.Epub 2017/04/13

4. Al-Goblan AS, Al-Alfi MA, Khan MZ. Mechanism linking diabetes mellitus and obesity. Diabetes Metab Syndr Obes. 2014; 7:587–91. Epub 2014/12/17. doi: 10.2147/DMSO.S67400.

5. Milic S, Lulic D, Stimac D. Non-alcoholic fatty liver disease and obesity: biochemical, metabolic and clinical presentations. World J Gastroenterol. 2014; 20(28):9330–7. Epub 2014/07/30. doi:10.3748/wjg.v20.i28.9330.

6. Lovren F, Teoh H, Verma S. Obesity and atherosclerosis: mechanistic insights. Can J Cardiol. 2015; 31(2):177–83. Epub 2015/02/11. doi:10.1016/j.cjca.2014.11.031.

7. Kachur S, Lavie CJ, de Schutter A, Milani RV, Ventura HO. Obesity and cardiovascular diseases. Minerva Med. 2017; 108(3):212–28. Epub 2017/02/06. doi:10.23736/S0026-4806.17.05022-4.

8. Basen-Engquist K, Chang M. Obesity and cancer risk: recent review and evidence. Curr Oncol Rep. 2011; 13(1):71–6. Epub 2010/11/17. doi:10.1007/s11912-010-0139-7.

9. Fock KM, Khoo J. Diet and exercise in management of obesity and overweight. J Gastroenterol Hepatol. 2013; 28 Suppl 4:59–63. doi:10.1111/jgh.12407.

10. Nguyen NT, Varela JE. Bariatric surgery for obesity and metabolic disorders: state of the art. Nat Rev Gastroenterol Hepatol. 2017; 14(3):160–9. doi:10.1038/nrgastro.2016.170.

11. Caterson ID. Medical management of obesity and its complications. Ann Acad Med Singapore. 2009; 38(1):22–7. Epub 2009/02/18.

12. Golden A. Current pharmacotherapies for obesity: A practical perspective. J Am Assoc Nurse Pract. 2017; 29(S1):S43–S52. doi:10.1002/2327-6924.12519.

13. Boutens L, Stienstra R. Adipose tissue macrophages: going off track during obesity. Diabetologia. 2016; 59(5):879–94. doi:10.1007/s00125-016-3904-9.

14. Coelho M, Oliveira T, Fernandes R. Biochemistry of adipose tissue: an endocrine organ. Arch Med Sci. 2013; 9(2):191–200. Epub 2013/05/15. doi:10.5114/aoms.2013.33181.

15. Khan M, Joseph F. Adipose tissue and adipokines: the association with and application of adipokines in obesity. Scientifica (Cairo). 2014; 328592. Epub 2014/10/14. doi:10.1155/2014/328592.

16. Choe SS, Huh JY, Hwang IJ, Kim JI, Kim JB. Adipose Tissue Remodeling: Its Role in Energy Metabolism and Metabolic Disorders. Front Endocrinol (Lausanne). 2016; 7:30. Epub 2016/05/06. doi: 10.3389/fendo.2016.00030.

17. Bartelt A, Heeren J. Adipose tissue browning and metabolic health. Nat Rev Endocrinol. 2014; 10(1):24–36. Epub 2013/10/23. doi: 10.1038/nrendo.2013.204.

18. Saely CH, Geiger K, Drexel H. Brown versus white adipose tissue: a mini-review. Gerontology. 2012; 58(1):15–23. Epub 2010/12/08. doi: 10.1159/000321319.

19. Sanchez-Gurmaches J, Hung CM, Guertin DA. Emerging Complexities in Adipocyte Origins and Identity. Trends Cell Biol. 2016; 26(5):313–26. Epub 2016/02/15. doi: 10.1016/j.tcb.2016.01.004.

20. Balistreri CR, Caruso C, Candore G. The role of adipose tissue and adipokines in obesity-related inflammatory diseases. Mediators Inflamm. 2010; 802078. doi: 10.1155/2010/802078.

21. Kim SH, Plutzky J. Brown Fat and Browning for the Treatment of Obesity and Related Metabolic Disorders. Diabetes Metab J. 2016; 40(1):12–21. Epub 2016/02/26. doi: 10.4093/dmj.2016.40.1.12.

22. Oakes ND, Kjellstedt A, Thalen P, Ljung B, Turner N. Roles of Fatty Acid oversupply and impaired oxidation in lipid accumulation in tissues of obese rats. J Lipids. 2013; 420754. Epub 2013/06/14. doi: 10.1155/2013/420754.

23. Suganami T, Tanaka M, Ogawa Y. Adipose tissue inflammation and ectopic lipid accumulation. Endocr J. 2012;59(10):849–57. Epub 2012/08/11.

24. Murano I, Barbatelli G, Parisani V, Latini C, Muzzonigro G, Castellucci M, et al. Dead adipocytes, detected as crown-like structures, are prevalent in visceral fat depots of genetically obese mice. J Lipid Res. 2008; 49(7):1562–8. Epub 2008/04/09. doi: 10.1194/jlr.M800019-JLR200.

25. Asghar A, Sheikh N. Role of immune cells in obesity induced low grade inflammation and insulin resistance. Cell Immunol. 2017; 315:18–26. Epub 2017/03/14. doi: 10.1016/j.cellimm.2017.03.001.

26. A.M. Castro LEM-dlC, C.A. Pantoja-Meléndez. Low-grade inflammation and its relation to obesity and chronic degenerative diseases. Rev Méd Hosp Gen Mex. 2017; 80(2):101–5.

27. Bell CJ, Sun Y, Nowak UM, Clark J, Howlett S, Pekalski ML, et al. Sustained in vivo signaling by long-lived IL-2 induces prolonged increases of regulatory T cells. J Autoimmun. 2015; 56:66–80. Epub 2014/12/03. doi:10.1016/j.jaut.2014.10.002.

28. Anderson PM, Sorenson MA. Effects of route and formulation on clinical pharmacokinetics of interleukin-2. Clin Pharmacokinet. 1994; 27(1):19–31. Epub 1994/07/01. doi: 10.2165/00003088-199427010-00003.

29. Lee SY, Cho ML, Oh HJ, Ryu JG, Park MJ, Jhun JY, et al. Interleukin-2/antiinterleukin-2 monoclonal antibody immune complex suppresses collagen-induced arthritis in mice by fortifying interleukin-2/STAT5 signalling pathways. Immunology. 2012; 137(4):305–16. Epub 2012/11/22. doi: 10.1111/imm.12008.

30. Wang YM, Alexander SI. IL-2/anti-IL-2 complex: a novel strategy of in vivo regulatory T cell expansion in renal injury. J Am Soc Nephrol. 2013; 24(10):1503–4. Epub 2013/08/21. doi: 10.1681/ASN.2013070718.

31. Webster KE, Walters S, Kohler RE, Mrkvan T, Boyman O, Surh CD, et al. In vivo expansion of T reg cells with IL-2-mAb complexes: induction of resistance to EAE and long-term acceptance of islet allografts without immunosuppression. J Exp Med. 2009; 206(4):751–60. Epub 2009/04/01. doi: 10.1084/jem.20082824.

32. Moon BI, Kim HR, Choi EJ, Kie JH, Seoh JY. Attenuation of collagen-induced arthritis by hyperbaric oxygen therapy through altering immune balance in favor of regulatory T cells. Undersea Hyperb Med. 2017; 44(4):321–30. Epub 2017/08/08.

33. Novak S, Drenjancevic I, Vukovic R, Kellermayer Z, Cosic A, Tolusic Levak M, et al. Anti-Inflammatory Effects of Hyperbaric Oxygenation during DSS-Induced Colitis in BALB/c Mice Include Changes in Gene Expression of HIF-1alpha, Proinflammatory Cytokines, and Antioxidative Enzymes. Mediators Inflamm. 2016; 7141430. Epub 2016/09/23. doi: 10.1155/2016/7141430.

34. Tuk B, Tong M, Fijneman EM, van Neck JW. Hyperbaric oxygen therapy to treat diabetes impaired wound healing in rats. PLoS One. 2014; 9(10):e108533. Epub 2014/10/21. doi: 10.1371/journal.pone.0108533.

35. Fukaya E, Hopf HW. HBO and gas embolism. Neurol Res. 2007; 29(2):142–5. Epub 2007/04/19. doi: 10.1179/016164107X174165.

36. Gurdol F, Cimsit M, Oner-Iyidogan Y, Korpinar S, Yalcinkaya S, Kocak H. Early and late effects of hyperbaric oxygen treatment on oxidative stress parameters in diabetic patients. Physiol Res. 2008; 57(1):41–7. Epub 2007/01/17.

37. Kim HR, Kim JH, Choi EJ, Lee YK, Kie JH, Jang MH, et al. Hyperoxygenation attenuated a murine model of atopic dermatitis through raising skin level of ROS. PLoS One. 2014; 9(10):e109297. Epub 2014/10/03. doi: 10.1371/journal.pone.0109297.

38. Kim HR, Lee A, Choi EJ, Hong MP, Kie JH, Lim W, et al. Reactive oxygen species prevent imiquimod-induced psoriatic dermatitis through enhancing regulatory T cell function. PLoS One. 2014; 9(3):e91146. Epub 2014/03/13. doi: 10.1371/journal.pone.0091146.

39. Tsuneyama K, Chen YC, Fujimoto M, Sasaki Y, Suzuki W, Shimada T, et al. Advantages and disadvantages of hyperbaric oxygen treatment in mice with obesity hyperlipidemia and steatohepatitis. ScientificWorldJournal. 2011; 11:2124–35. doi: 10.1100/2011/380236.

40. Wilkinson D, Chapman IM, Heilbronn LK. Hyperbaric oxygen therapy improves peripheral insulin sensitivity in humans. Diabet Med. 2012; 29(8):986–9. doi: 10.1111/j.1464-5491.2012.03587.x.

41. Wilkinson D, Nolting M, Mahadi MK, Chapman I, Heilbronn L. Hyperbaric oxygen therapy increases insulin sensitivity in overweight men with and without type 2 diabetes. Diving Hyperb Med. 2015; 45(1):30–6.

42. Takahashi Y, Fukusato T. Histopathology of nonalcoholic fatty liver disease/nonalcoholic steatohepatitis. World J Gastroenterol. 2014; 20(42):15539–48. Epub 2014/11/18. doi: 10.3748/wjg.v20.i42.15539.

43. Halberg N, et al.,. Hypoxia-inducible factor 1 alpha induces fibrosis and insulin resistance in white adipose tissue. Mol cell biol. 2009; 29(16):4467–83.

44. Takikawa A, Mahmood A, Nawaz A, Kado T, Okabe K, Yamamoto S, et al. HIF-1alpha in Myeloid Cells Promotes Adipose Tissue Remodeling Toward Insulin Resistance. Diabetes. 2016; 65(12):3649–59. Epub 2016/09/15. doi: 10.2337/db16-0012.

45. Sun K, Halberg N, Khan M, Magalang UJ, Scherer PE. Selective inhibition of hypoxia-inducible factor 1alpha ameliorates adipose tissue dysfunction. Mol Cell Biol. 2013; 33(5):904–17. Epub 2012/12/20. doi: 10.1128/MCB.00951-12.

46. Mylonis I, Sembongi H, Befani C, Liakos P, Siniossoglou S, Simos G. Hypoxia causes triglyceride accumulation by HIF-1-mediated stimulation of lipin 1 expression. J Cell Sci. 2012; 125(Pt 14):3485–93. Epub 2012/04/03. doi: 10.1242/jcs.106682.

47. Norouzirad R, Gonzalez-Muniesa P, Ghasemi A. Hypoxia in Obesity and Diabetes: Potential Therapeutic Effects of Hyperoxia and Nitrate. Oxid Med Cell Longev. 2017; 5350267. Epub 2017/06/14. doi: 10.1155/2017/5350267.

48. Patel P AN. Role of subcutaneous adipose tissue in the pathogenesis of insulin resistance. J Obes 2013; 489187. doi: 10.1155/2013/489187.

49. Stanford KI MR, Townsend KL, An D, Nygaard EB, Hitchcox KM, Markan KR, Nakano K, Hirshman MF, Tseng YH, Goodyear LJ. Brown adipose tissue regulates glucose homeostasis and insulin sensitivity. J Clin Invest. 2013; 123(1):215–23. doi: 10.1172/JCI62308.

50. Chondronikola M, Volpi E, Borsheim E, Porter C, Annamalai P, Enerback S, et al. Brown adipose tissue improves whole-body glucose homeostasis and insulin sensitivity in humans. Diabetes. 2014; 63(12):4089–99. Epub 2014/07/25. doi: 10.2337/db14-0746.

